# Ancient human genomes suggest three ancestral populations for present-day Europeans

**DOI:** 10.1101/001552

**Authors:** Iosif Lazaridis, Nick Patterson, Alissa Mittnik, Gabriel Renaud, Swapan Mallick, Karola Kirsanow, Peter H. Sudmant, Joshua G. Schraiber, Sergi Castellano, Mark Lipson, Bonnie Berger, Christos Economou, Ruth Bollongino, Qiaomei Fu, Kirsten I. Bos, Susanne Nordenfelt, Heng Li, Cesare de Filippo, Kay Prüfer, Susanna Sawyer, Cosimo Posth, Wolfgang Haak, Fredrik Hallgren, Elin Fornander, Nadin Rohland, Dominique Delsate, Michael Francken, Jean-Michel Guinet, Joachim Wahl, George Ayodo, Hamza A. Babiker, Graciela Bailliet, Elena Balanovska, Oleg Balanovsky, Ramiro Barrantes, Gabriel Bedoya, Haim Ben-Ami, Judit Bene, Fouad Berrada, Claudio M. Bravi, Francesca Brisighelli, George Busby, Francesco Cali, Mikhail Churnosov, David E. C. Cole, Daniel Corach, Larissa Damba, George van Driem, Stanislav Dryomov, Jean-Michel Dugoujon, Sardana A. Fedorova, Irene Gallego Romero, Marina Gubina, Michael Hammer, Brenna Henn, Tor Hervig, Ugur Hodoglugil, Aashish R. Jha, Sena Karachanak-Yankova, Rita Khusainova, Elza Khusnutdinova, Rick Kittles, Toomas Kivisild, William Klitz, Vaidutis Kučinskas, Alena Kushniarevich, Leila Laredj, Sergey Litvinov, Theologos Loukidis, Robert W. Mahley, Béla Melegh, Ene Metspalu, Julio Molina, Joanna Mountain, Klemetti Näkkäläjärvi, Desislava Nesheva, Thomas Nyambo, Ludmila Osipova, Jüri Parik, Fedor Platonov, Olga Posukh, Valentino Romano, Francisco Rothhammer, Igor Rudan, Ruslan Ruizbakiev, Hovhannes Sahakyan, Antti Sajantila, Antonio Salas, Elena B. Starikovskaya, Ayele Tarekegn, Draga Toncheva, Shahlo Turdikulova, Ingrida Uktveryte, Olga Utevska, René Vasquez, Mercedes Villena, Mikhail Voevoda, Cheryl Winkler, Levon Yepiskoposyan, Pierre Zalloua, Tatijana Zemunik, Alan Cooper, Cristian Capelli, Mark G. Thomas, Andres Ruiz-Linares, Sarah A. Tishkoff, Lalji Singh, Kumarasamy Thangaraj, Richard Villems, David Comas, Rem Sukernik, Mait Metspalu, Matthias Meyer, Evan E. Eichler, Joachim Burger, Montgomery Slatkin, Svante Pääbo, Janet Kelso, David Reich, Johannes Krause

## Abstract

We sequenced genomes from a ∼7,000 year old early farmer from Stuttgart in Germany, an ∼8,000 year old hunter-gatherer from Luxembourg, and seven ∼8,000 year old hunter-gatherers from southern Sweden. We analyzed these data together with other ancient genomes and 2,345 contemporary humans to show that the great majority of present-day Europeans derive from at least three highly differentiated populations: West European Hunter-Gatherers (WHG), who contributed ancestry to all Europeans but not to Near Easterners; Ancient North Eurasians (ANE), who were most closely related to Upper Paleolithic Siberians and contributed to both Europeans and Near Easterners; and Early European Farmers (EEF), who were mainly of Near Eastern origin but also harbored WHG-related ancestry. We model these populations’ deep relationships and show that EEF had ∼44% ancestry from a “Basal Eurasian” lineage that split prior to the diversification of all other non-African lineages.

---

Ancient DNA studies have demonstrated that migration played a major role in the introduction of agriculture to Europe, as early farmers were genetically distinct from hunter-gatherers^1,2^ and closer to present-day Near Easterners^2,3^. Modelling the ancestry of present-day Europeans as a simple mixture of two ancestral populations^2^, however, does not take into account their genetic affinity to an Ancient North Eurasian (ANE) population^4,5^ who also contributed genetically to Native Americans^6^. To better understand the deep ancestry of present-day Europeans, we sequenced nine ancient genomes that span the transition from hunting and gathering to agriculture in Europe (Fig. 1A; Extended Data Fig. 1): “Stuttgart” (19-fold coverage), a ∼7,000 year old skeleton found in Germany in the context of artifacts from the first widespread Neolithic farming culture of central Europe, the *Linearbandkeramik*; “Loschbour” (22-fold coverage), an ∼8,000 year old skeleton from the Loschbour rock shelter in Heffingen, Luxembourg, discovered in the context of Mesolithic hunter-gatherer artifacts (SI1; SI2); and seven samples (0.01-2.4-fold coverage) from an ∼8,000 year old Mesolithic hunter-gatherer burial in Motala, Sweden.

**Figure 1:**
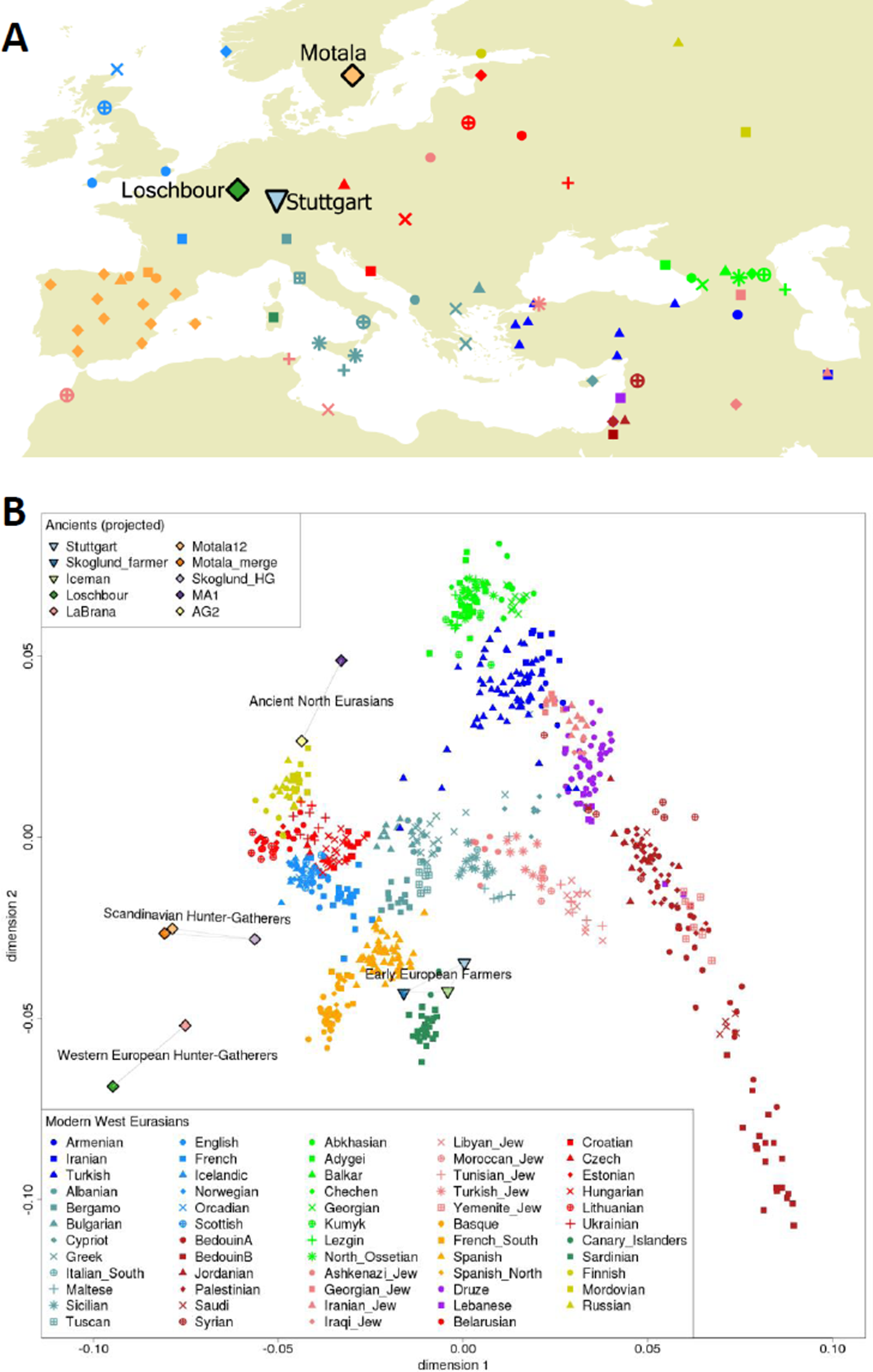
Map of West Eurasian populations and Principal Component Analysis. (a) Geographical locations of ancient and present-day samples, with color coding matching the PCA. We show all sampling locations for each population, which results in multiple points for some populations (e.g., Spain). (b) PCA on all present-day West Eurasians, with the ancient and selected eastern non-Africans projected. European hunter-gatherers fall beyond present-day Europeans in the direction of European differentiation from the Near East. Stuttgart clusters with other Neolithic Europeans and present-day Sardinians. MA1 falls outside the variation of present-day West Eurasians in the direction of southern-northern differentiation along dimension 2 and between the European and Near Eastern clines along dimension 1.

A central challenge is to show that DNA sequences retrieved from ancient samples are authentic and not due to present-day human contamination. The rate of C→T and G→A mismatches to the human genome at the ends of the molecules in libraries from each of the ancient samples exceeds 20%, a signature that suggests the DNA is largely ancient^7,8^ (SI3). We inferred mitochondrial DNA (mtDNA) consensus sequences, and based on the number of sites that differed, estimated contamination rates of 0.3% for Loschbour, 0.4% for Stuttgart, and 0.01%-5% for the Motala individuals (SI3). We inferred similar levels of contamination for the nuclear DNA of Loschbour (0.4%) and Stuttgart (0.3%) using a maximum-likelihood-based test (SI3). The effective contamination rate for the high coverage samples is likely to be far lower, as consensus diploid genotype calling (SI2) tends to reduce the effects of a small fraction of contaminating reads.

Stuttgart belongs to mtDNA haplogroup T2, typical of Neolithic Europeans^9^, while Loschbour and all Motala individuals belong to haplogroups U5 and U2, typical of pre-agricultural Europeans^1,7^ (SI4). Based on the ratio of reads aligning to chromosomes X and Y, Stuttgart is female, while Loschbour and five of seven Motala individuals are male^10^ (SI5). Loschbour and the four Motala males whose haplogroups we could determine all belong to Y-chromosome haplogroup I, suggesting that this was a predominant haplogroup in pre-agricultural northern Europeans analogous to mtDNA haplogroup U^11^ (SI5).

We carried out most of our sequencing on libraries prepared in the presence of uracil DNA glycosylase (UDG), which reduces C→T and G→A errors due to ancient DNA damage (SI3). We first confirm that the ancient samples had statistically indistinguishable levels of Neandertal ancestry to each other (∼2%) and to present-day Eurasians (SI6), and so we do not consider this further in our analyses of population relationships. We report analyses that leverage the type of information that can only be obtained from deep coverage genomes, mostly focusing on Loschbour and Stuttgart, and for some analyses also including Motala12 (2.4×) and La Braña from Mesolithic Iberia (3.4×)^12^. Heterozygosity, the number of differences per nucleotide between an individual’s two chromosomes, is 0.00074 for Stuttgart, at the high end of present-day Europeans, and 0.00048 for Loschbour, lower than in any present-day humans (SI2). Through comparison of Loschbour’s two chromosomes we find that this low diversity is not due to recent inbreeding but instead due to a population bottleneck in this individual’s more distant ancestors (Extended Data Fig. 2). Regarding alleles that affect phenotype, we find that the *AMY1* gene coding for salivary amylase had 5, 6, 13, and 16 copies in La Braña^12^, Motala12, Loschbour and Stuttgart respectively; these numbers are within the range of present-day Europeans (SI7), suggesting that high copy counts of *AMY1* are not entirely due to selection since the switch to agriculture^13^. The genotypes at SNPs associated with lactase persistence indicate that Stuttgart, Loschbour, and Motala12 were unable to digest milk as adults. Both Loschbour and Stuttgart likely had dark hair (>99% probability); Loschbour, like La Braña and Motala12, likely had blue or intermediate-colored eyes (>75% probability), while Stuttgart most likely had brown eyes (>99% probability) (SI8). Neither Loschbour nor La Braña carries the skin-lightening allele in *SLC24A5* that is homozygous in Stuttgart and nearly fixed in Europeans today, indicating that they probably had darker skin^12^. However, Motala12 carries at least one copy of the derived allele, indicating that this locus was already polymorphic in Europeans prior to the advent of agriculture.

To place the ancient European genomes in the context of present-day human genetic variation, we assembled a dataset of 2,345 present-day humans from 203 populations genotyped at 594,924 autosomal single nucleotide polymorphisms (SNPs)^5^ (SI9) (Extended Data Table 1). We used ADMIXTURE^14^ to identify 59 “West Eurasian” populations (777 individuals) that cluster with Europe and the Near East (SI9 and Extended Data Fig. 3). Principal component analysis (PCA)^15^ (SI10) (Fig. 1B) reveals a discontinuity between the Near East and Europe, with each showing north-south clines bridged only by a few populations of mainly Mediterranean origin. Our PCA differs from previous studies that showed a correlation with the map of Europe^16,17^, which we determined is due to our study having relatively fewer central and northwestern Europeans, and more Near Easterners and eastern Europeans (SI10). We projected^18^ the newly sequenced and previously published^2,6,12,19^ ancient genomes onto the first two PCs inferred from present-day samples (Fig. 1B). MA1 and AG2, both Upper Paleolithic hunter-gatherers from Lake Baikal^6^ in Siberia, project at the northern end of the PCA, suggesting an “Ancient North Eurasian” meta-population (ANE). European hunter-gatherers from Spain, Luxembourg, and Sweden fall outside the genetic variation of West Eurasians in the direction of European differentiation from the Near East, with a “West European Hunter-Gatherer” (WHG) cluster including Loschbour and La Braña^12^, and a “Scandinavian Hunter-Gatherer” (SHG) cluster including the Motala individuals and ∼5,000 year old hunter-gatherers from the Swedish Pitted Ware Culture^2^. An “Early European Farmer” (EEF) cluster includes Stuttgart, the ∼5,300 year old Tyrolean Iceman^19^ and a ∼5,000 year old southern Swedish farmer^2^, and is near present-day Sardinians^2,19^.

PCA gradients of genetic variation may arise under very different histories^20^. To test if they reflect population mixture events or are entirely due to genetic drift within West Eurasia, we computed an *f_4_-*statistic^18^ that tests whether the ancient MA1 from Siberia shares more alleles with a *Test* West Eurasian population or with Stuttgart. We find that *f_4_(Test, Stuttgart; MA1, Chimp)* is positive for many West Eurasians, which must be due to variable degrees of admixture with ancient populations related to MA1 (Extended Data Fig. 4). We also find that *f_4_(Test, Stuttgart; Loschbour, Chimp)* is nearly always positive in Europeans and always negative in Near Easterners, indicating that Europeans have more ancestry from ancient populations related to Loschbour than do Near Easterners (Extended Data Fig. 4). To investigate systematically the history of population mixture in West Eurasia, we computed all possible statistics of the form *f_3_(X; Ref_1_, Ref_2_)* (SI11). An *f_3_*-statistic is expected to be positive if no admixture has taken place, but if *X* is admixed between populations related to *Ref_1_* and *Ref_2_*, it can be negative^5^. We tested all possible pairs of *Ref_1_*, *Ref_2_* chosen from the list of 192 present-day populations with at least four individuals, and five ancient genomes (SI11). The lowest *f_3_*-statistics for Europeans are usually negative (93% are >4 standard errors below zero using a standard error from a block jackknife^5,21^). The most negative statistic (Table 1) always involves at least one ancient individual as a reference, and for Europeans it is nearly always significantly lower than the most negative statistic obtained using only present-day populations as references (SI11). MA1 is a better surrogate (Extended Data Fig. 5) for Ancient North Eurasian ancestry than the Native American Karitiana who were first used to represent this component of ancestry in Europe^4,5^. Motala12 never appears as one of the references, suggesting that SHG may not be a source for Europeans. Instead, present-day European populations usually have their lowest *f_3_* with either the (EEF, ANE) or (WHG, Near East) pair (SI11, Extended Data Table 1). For Near Easterners, the lowest *f_3_*-statistic always takes as references Stuttgart and a population from Africa, the Americas, South Asia, or MA1 (Table 1), reflecting the fact that both Stuttgart and present-day Near Easterners harbor ancestry from ancient Near Easterners. Extended Data Fig. 6 plots statistics of the form *f_4_(West Eurasian X, Chimp; Ancient_1_, Ancient_2_)* onto a map, showing strong gradients in the relatedness to Stuttgart (EEF), Loschbour (WHG) and MA1 (ANE).

**Table 1:** Lowest *f_3_*-statistics for each West Eurasian population

| *Ref<sub>1</sub>* | *Ref<sub>2</sub>* | Target for which these two references give the lowest $f_3(X; Ref_1, Ref_2)$ |
| --- | --- | --- |
| **WHG** | **EEF** | Sardinian<sup>***</sup> |
| **WHG** | **Near East** | Basque, Belarusian, Czech, English, Estonian, Finnish, French_South, Icelandic, Lithuanian, Mordovian, Norwegian, Orcadian, Scottish, Spanish, Spanish_North, Ukrainian |
| **EEF** | **ANE** | Abkhasian<sup>***</sup>, Albanian, Ashkenazi_Jew<sup>****</sup>, Bergamo, Bulgarian, Chechen<sup>****</sup>, Croatian, Cypriot<sup>****</sup>, Druze<sup>**</sup>, French, Greek, Hungarian, Lezgin, MA1, Maltese, Sicilian, Turkish_Jew, Tuscan |
| **EEF** | **Native American** | Adygei, Balkar, Iranian, Kumyk, North_Ossetian, Turkish |
| **EEF** | **African** | BedouinA, BedouinB†, Jordanian, Lebanese, Libyan_Jew, Moroccan_Jew, Palestinian, Saudi<sup>****</sup>, Syrian, Tunisian_Jew<sup>***</sup>, Yemenite_Jew<sup>***</sup> |
| **EEF** | **South Asian** | Armenian, Georgian<sup>****</sup>, Georgian_Jew<sup>*</sup>, Iranian_Jew<sup>***</sup>, Iraqi_Jew<sup>***</sup> |

Note: WHG = Loschbour or LaBraña; EEF=Stuttgart; ANE=MA1; Native American=Piapoco; African=Esan, Gambian, or Kgalagadi; South Asian=GujaratiC or Vishwabrahmin. Statistics are negative with $Z < -4$ unless otherwise noted: † (positive) or \*, \*\*, \*\*\*, \*\*\*\*, to indicate Z less than 0, -1, -2, and -3 respectively. The complete list of statistics can be found in Extended Data Table 1.

We determined formally that a minimum of three source ancestral populations are needed to explain the data for many European populations taken together by studying the correlation patterns of all possible statistics of the form *f_4_(Test_base_, Test_i_; O_base_, O_j_)* (SI12). Here *Test*_*base*_ is a reference European population and *Test*_*i*_ the set of all other European *Test* populations; *O*_*base*_ is a reference outgroup population, and *O_i_* the set of other outgroups (ancient DNA samples, Onge, Karitiana, and Mbuti). The rank of the *(i, j)* matrix reflects the minimum number of source populations that contributed to the *Test* populations^22,23^. For a pool of 23 *Test* populations comprising most present-day Europeans, this analysis rejects descent from just two sources (P<10^-12^ by a Hotelling T-test^23^). However, three source populations are consistent with the data after excluding the Spanish who have evidence for African admixture^24–26^ (P=0.019, not significant after multiple-hypothesis correction). Our finding of at least three source populations is also qualitatively consistent with the results from ADMIXTURE (SI9), PCA (Fig. 1B, SI10) and *f*-statistics (Extended Data Table 1, Extended Data Fig. 6, SI11, SI12). We caution that the finding of three sources could be consistent with a larger number of mixture events, as the method cannot distinguish between one or more mixture events if they are from the same set of sources. Our analysis also does not assume that the inferred source populations were themselves unadmixed; indeed, the positive *f_4_(Stuttgart, X; Loschbour, Chimp)* statistics obtained when *X* is a Near Eastern population (Extended Data Table 1) implies that EEF had some WHG-related ancestry, which we show in SI13 was at least 0% and less than 45%.

Motivated by the evidence of at least three source populations for present-day Europeans, we set out to develop a model consistent with our data. To constrain our search space for modeling, we first studied *f_4_-*statistics comparing the ancient individuals from Europe and Siberia and diverse eastern non-African groups (Oceanians, East Asians, Siberians, Native Americans, and Onge from the Andaman Islands^27^) (SI14). We find that: (1) Loschbour (WHG) and Stuttgart (EEF) share more alleles with each other than either does with MA1 (ANE), as might be expected by geography, but MA1 shares more alleles with Loschbour than with Stuttgart, indicating a link between Eurasian hunter-gatherers to the exclusion of European farmers; (2) Eastern non-Africans share more alleles with Eurasian hunter-gatherers (MA1, Loschbour, La Braña, and Motala12) than with Stuttgart; (3) Every eastern non-African population except for Native Americans and Siberians is equally closely related to diverse Eurasian hunter-gatherers, but Native Americans and Siberians share more alleles with MA1 than with European hunter-gatherers; and (4) Eurasian hunter-gatherers and Stuttgart both share more alleles with Native Americans than with other eastern non-Africans. We use the ADMIXTUREGRAPH^18^ software to search for a model of population relationships (a tree structure augmented by admixture events) that is consistent with these observations. We explored models with 0, 1, or 2 admixture events in the ancestry of the three ancient source populations and eastern non-Africans, and identified a single model with two admixture events that fit the data. The successful model (Fig. 2A) includes the previously reported gene flow into Native Americans from an MA1-like population^6^, as well as the novel inference that Stuttgart is partially (44 ± 10%) derived from a “Basal Eurasian” lineage that split prior to the separation of eastern non-Africans from the common ancestor of WHG and ANE. If this model is accurate, the ANE/WHG split must have occurred >24,000 years ago since this is the age^6^ of MA1 and this individual is on the ANE lineage. The WHG must then have split from eastern non-Africans >40,000 years ago, as this is the age of the Chinese Tianyuan sample which clusters with eastern non-Africans to the exclusion of Europeans^28^. The Basal Eurasian split would then have to be even older. A Basal Eurasian lineage in the Near East is plausible given the presence of anatomically modern humans in the Levant^29^ ∼100 thousand years ago and African-related tools likely made by modern humans in Arabia^30,31^. Alternatively, evidence for gene flow between the Near East and Africa^32^, and African morphology in pre-farming Natufians^33^ from Israel, may also be consistent with the population representing a later movement of humans out of Africa and into the Near East.

**Figure 2:**
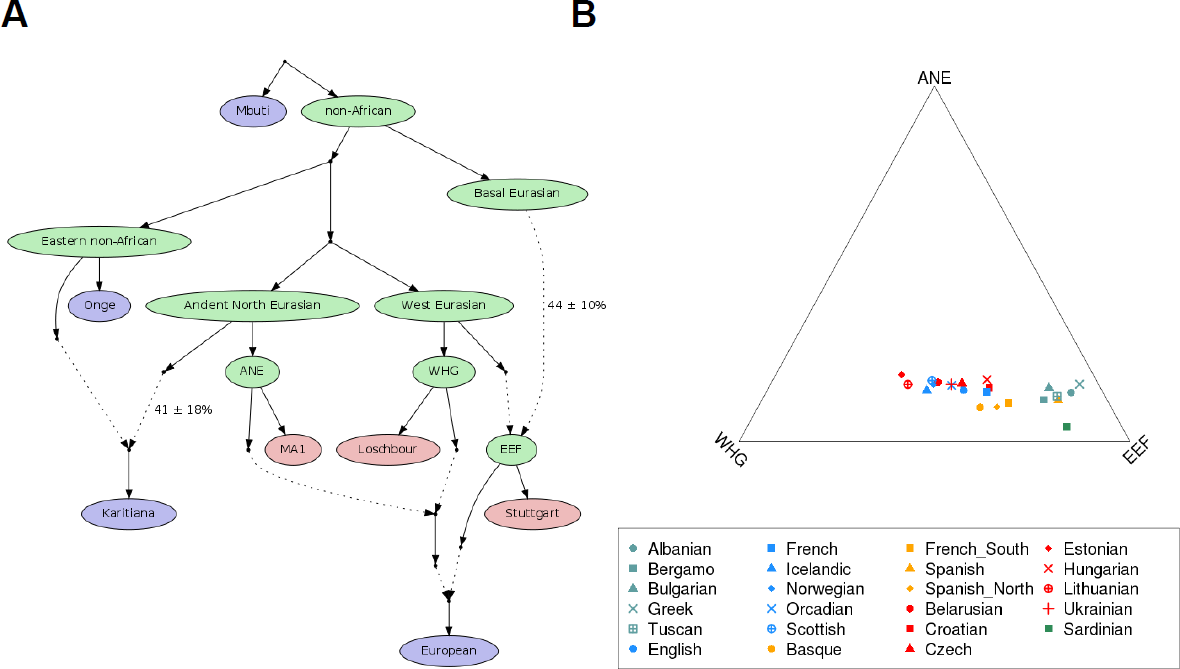
Modeling of West Eurasian population history. (a) A three-way mixture model that is a statistical fit to the data for many European populations, ancient DNA samples, and non-European populations. Present-day samples are colored in blue, ancient samples in red, and reconstructed ancestral populations in green. Solid lines represent descent without mixture, and dashed lines represent admixture events. For the two mixture events relating the highly divergent ancestral populations, we print estimates for the mixture proportions as well as one standard error. (b) We plot the proportions of ancestry from each of three inferred ancestral populations (EEF, ANE and WHG) as inferred from the model-based analysis.

We tested the robustness of the ADMIXTUREGRAPH model in various ways. First, we verified that Stuttgart and the Iceman (EEF), and Loschbour and LaBraña (WHG) can be formally fit as clades (SI14). We also used the unsupervised MixMapper^4^ (SI15) and TreeMix^34^ software (SI16) to fit graph models; both found all the same admixture events. The statistics supporting our key inferences about history also provide consistent results when restricted to transversions polymorphisms not affected by ancient DNA damage, and when repeated with whole-genome sequencing data that is not affected by SNP ascertainment bias^35^ (Extended Data Table 2).

We next fit present-day European populations into our working model. We found that few European populations could be fit as 2-way mixtures, but nearly all were compatible with being 3-way mixtures of ANE/EEF/WHG (SI14). Mixture proportions (Fig. 2B; Extended Data Table 3) inferred via our model are consistent with those from an independent method that relates European populations to diverse outgroups using *f_4_*-statistics while making much weaker modeling assumptions (only assuming that MA1 is an unmixed descendent of ANE, Loschbour of WHG, and Stuttgart of EEF; SI17). These analyses allow us to infer that EEF ancestry in Europe today ranges from ∼30% in the Baltic region to ∼90% in the Mediterranean, a gradient that is also consistent with patterns of identity-by-descent (IBD) sharing^36^ (SI18) and chromosome painting^37^ (SI19) in which Loschbour shares more segments with northern Europeans and Stuttgart with southern Europeans. Our estimates suggest that Southern Europeans inherited their European hunter-gatherer ancestry mostly via EEF ancestors (Extended Data Fig. 6), while Northern Europeans acquired up to 50% additional WHG ancestry. Europeans have a larger proportion of WHG than ANE ancestry (WHG/(WHG+ANE) = 0.6-0.8) with the ANE ancestry never being larger than ∼20%. (By contrast, in the Near East there is no detectible WHG ancestry, but substantial ANE ancestry, up to ∼29% in the North Caucasus) (SI14). While ANE ancestry was not as pervasive in Europe during the agricultural transition as it is today (we do not detect it in either Loschbour or Stuttgart), it was already present, since MA1 shares more alleles with Motala12 (SHG) than with Loschbour, and Motala12 fits as a mixture of 81% WHG and 19% ANE (SI14).

Two sets of European populations are poor fits. Sicilians, Maltese, and Ashkenazi Jews have EEF estimates beyond the 0-100% interval (SI17) and cannot be jointly fit with other Europeans (SI14). These populations may have more Near Eastern ancestry than can be explained via EEF admixture (SI14), consistent with their falling in the gap between European and Near Eastern populations in Fig. 1B. Finns, Mordovians and Russians from northeastern Europe also do not fit (SI14; Extended Data Table 3). To better understand this, we plotted *f_4_(X, Bedouin2; Han, Mbuti)* against *f_4_(X, Bedouin2; MA1, Mbuti).* These statistics measure the degree of a European population’s allele sharing with Han Chinese or MA1 (Extended Data Fig. 7). Europeans fall on a line of slope >1 in the plot of these two statistics. However, northeastern Europeans including Chuvash and Saami (which we add in to the analysis) fall away from this line in the direction of East Asians. This is consistent with East Asian (most likely Siberian) gene flow into northeastern Europeans, some of which may be more recent^38^ than the original ANE admixture (SI14).

Three questions seem particularly important to address in follow-up work. Where did the EEF obtain their WHG ancestry? Southeastern Europe is a candidate as it lies along the path from Anatolia into central Europe^39^. When and where the ancestors of present-day Europeans first acquire their ANE ancestry? Based on discontinuity in mtDNA haplogroup frequencies, this may have occurred ∼5,500-4,000 years ago^40^ in Central Europe. When and where did Basal Eurasians mix into the ancestors of the EEF? An important aim for future work should be to collect DNA from additional ancient samples to illuminate these transformations.

## Methods Summary

We extracted DNA from nine sets of ancient human remains and converted the extracts into Illumina sequencing libraries in dedicated clean rooms. We assessed whether sequences for these libraries were consistent with genuine ancient DNA by searching for characteristic deaminations at the ends of molecules^7,8^. We also tested for contamination by searching for evidence of mixture of DNA from multiple individuals. For large-scale shotgun sequencing we used libraries that we made in the presence of the enzymes Uracil-DNA-glycosylase and endonuclease VIII, which reduce the rate of ancient DNA-induced errors. After removal of duplicated molecules, we called consensus genotypes for the high coverage samples using the Genome Analysis Toolkit^41^. We merged the data with published ancient genomes, as well as with 2,345 present-day humans from 203 populations genotyped at 594,924 autosomal single nucleotide polymorphisms. We visualized population structure using Principal Component Analysis^15^ and ADMIXTURE^14^. To make inferences about population history, we used methods that can analyze allele frequency correlation statistics to detect population mixture^5^; that can estimate mixture proportions in the absence of accurate ancestral populations; that can infer the minimum number of source populations for a collection of tests population^23^; and that can assess formally the fit of genetic data to models of population history^5^.

**Supplementary Information** is linked to the online version of the paper. The fully public version of the Human Origins dataset can be found at http://genetics.med.harvard.edu/reichlab/Reich_Lab/Datasets.html. The full version of the dataset (including additional samples) is available to researchers who send a signed letter to DR indicating that they will abide by specified usage conditions.

## Acknowledgments

We are grateful to Cynthia Beall, Neil Bradman, Amha Gebremedhin, Damian Labuda, Maria Nelis and Anna Di Rienzo for sharing DNA samples; to Detlef Weigel, Christa Lanz, Verena Schünemann, Peter Bauer and Olaf Riess for support and access to DNA sequencing facilities; to Philip Johnson for advice on contamination estimation; and to Pontus Skoglund for sharing the graphics software that we used to generate Extended Data Fig. 6. We thank Kenneth Nordtvedt for alerting us about the existence of newly discovered Y-chromosome SNPs. The collections and methods for the Population Reference Sample (POPRES) discussed in SI18 are described in ref.^42^, and the dataset used for our analyses was obtained from dbGaP at http://www.ncbi.nlm.nih.gov/projects/gap/cgi-bin/study.cgi?study_id=phs000145.v4.p2 through dbGaP accession number phs000145.v1.p2. We thank all the volunteers who donated DNA; the staff of the Unità Operativa Complessa di Medicina Trasfusionale, Azienda Ospedaliera Umberto I, Siracusa, Italy for assistance in sample collection; and The National Laboratory for the Genetics of Israeli Populations for facilitating access to DNA. We thank colleagues at the Applied Genomics at the Children’s Hospital of Philadelphia, especially Hakon Hakonarson, Cecilia Kim, Kelly Thomas, and Cuiping Hou, for genotyping samples on the Human Origins array. JK is grateful for support from DFG grant # KR 4015/1-1, the Carl-Zeiss Foundation and the Baden Württemberg Foundation. SP acknowledges support from the Presidential Innovation Fund of the Max Planck Society. JGS acknowledges use of the Extreme Science and Engineering Discovery Environment (XSEDE), which is supported by NSF grant number OCI-1053575. EB and OB were supported by RFBR grants 13-06-00670, 13-04-01711, 13-04-90420 and by the Molecular and Cell Biology Program of the Presidium, Russian Academy of Sciences. BM was supported by grants OTKA 73430 and 103983. ASaj was supported by a Finnish Professorpool (Paulo Foundation) Grant. The Lithuanian sampling was supported by the LITGEN project (VP1-3.1-ŠMM-07-K-01-013), funded by the European Social Fund under the Global Grant Measure. AS was supported by Spanish grants SAF2008-02971 and EM 2012/045. OU was supported by Ukrainian SFFS grant F53.4/071. SAT was supported by NIH Pioneer Award 8DP1ES022577-04 and NSF HOMINID award BCS-0827436. KT was supported by an Indian CSIR Network Project (GENESIS: BSC0121). LS was supported by an Indian CSIR Bhatnagar Fellowship. RV, MM, JP and EM were supported by the European Union Regional Development Fund through the Centre of Excellence in Genomics to the Estonian Biocentre and University of Tartu and by a Estonian Basic Research grant SF0270177As08. MM was additionally supported by Estonian Science Foundation grant #8973. JGS and MS were supported by NIH grant GM40282. PHS and EEE were supported by NIH grants HG004120 and HG002385. DR and NP were supported by NSF HOMINID award BCS-1032255 and NIH grant GM100233. DR and EEE are Howard Hughes Medical Institute investigators.

## Author contributions

BB, EEE, JBu, MS, SP, JKe, DR and JKr supervised the study. IL, NP, AM, GR, SM, KK, PHS, JGS, SC, ML, QF, HL, CdF, KP, WH, MMey and DR analyzed genetic data. FH, EF, DD, MF, J-MG, JW, AC and JKr obtained human remains. AM, CE, RBo, KB, SS, CP, NR and JKr processed ancient DNA. IL, NP, SN, NR, GA, HAB, GBa, EB, OB, RBa, GBe, HB-A, JBe, FBe, CMB, FBr, GBJB, FC, MC, DECC, DCor, LD, GvD, SD, J-MD, SAF, IGR, MG, MH, BH, TH, UH, ARJ, SK-Y, RKh, EK, RKi, TK, WK, VK, AK, LL, SL, TL, RWM, BM, EM, JMol, JMou, KN, DN, TN, LO, JP, FP, OLP, VR, FR, IR, RR, HS, ASaj, ASal, EBS, ATar, DT, ST, IU, OU, RVa, MVi, MVo, CW, LY, PZ, TZ, CC, MGT, AR-L, SAT, LS, KT, RVi, DCom, RS, MMet, SP and DR assembled the genotyping dataset. IL, NP, DR and JKr wrote the manuscript with help from all co-authors.

## Author information

The aligned sequences are available through the Sequence Read Archive (SRA) under accession numbers that will be made available upon publication. The authors declare competing financial interests: UH is an employee of Illumina, TL is an employee of AMGEN, and JM is an employee of 23andMe. Correspondence and requests for materials should be addressed to David Reich or Johannes Krause.

## Methods

### Archeological context, sampling and DNA extraction

The Loschbour sample stems from a male skeleton excavated in 1935 at the Loschbour rock shelter in Heffingen, Luxembourg. The skeleton was AMS radiocarbon dated to 7,205 ± 50 years before present (OxA-7738; 6,220-5,990 cal BC)^43^. At the Palaeogenetics Laboratory in Mainz, material for DNA extraction was sampled from a molar (M48) after irradiation with UV-light, surface removal, and pulverization in a mixer mill. DNA extraction took place in the palaeogenetics facilities in the Institute for Archaeological Sciences at the University of Tübingen. Three extracts were made in total, one from 80 mg of powder using an established silica based protocol^44^ and two additional extracts from 90 mg of powder each with a protocol optimized for the recovery of short DNA molecules^45^.

The Stuttgart sample was taken from a female skeleton excavated in 1982 at the site Viesenhäuser Hof, Stuttgart-Mühlhausen, Germany. It was attributed to the Linearbandkeramik (5,500-4,800 BC) through associated pottery artifacts and the chronology was corroborated by radiocarbon dating of the stratigraphy^46^. Both sampling and DNA extraction took place in the Institute for Archaeological Sciences at the University of Tübingen. The M47 molar was removed and material from the inner part was sampled with a sterile dentistry drill. An extract was made using 40 mg of bone powder^45^.

The Motala individuals were recovered from the site of Kanaljorden in the town of Motala, Östergötland, Sweden, excavated between 2009 and 2013. The human remains at this site are represented by several adult skulls and one infant skeleton. All individuals are part of a ritual deposition at the bottom of a small lake. Direct radiocarbon dates on the remains range between 7,013 ± 76 and 6,701 ± 64 BP (6,361-5,516 cal BC), corresponding to the late Middle Mesolithic of Scandinavia. Samples were taken from the teeth of the nine best preserved skulls, as well as a femur and tibia. Bone powder was removed from the inner parts of the teeth or bones with a sterile dentistry drill. DNA from 100 mg of bone powder was extracted^47^ in the ancient DNA laboratory of the Archaeological Research Laboratory, Stockholm.

### Library preparation

Illumina sequencing libraries were prepared using either double- or single-stranded library preparation protocols^48,49^ (SI1). For high-coverage shotgun sequencing libraries, a DNA repair step with Uracil-DNA-glycosylase (UDG) and endonuclease VIII (endo VIII) treatment was included in order to remove uracil residues^50^. Size fractionation on a PAGE gel was also performed in order to remove longer DNA molecules that are more likely to be contaminants^49^. Positive and blank controls were carried along during every step of library preparation.

### Shotgun sequencing and read processing

All non-UDG-treated libraries were sequenced either on an Illumina Genome Analyzer IIx with 2×76 + 7 cycles for the Loschbour and Motala libraries, or on an Illumina MiSeq with 2×150 + 8 + 8 cycles for the Stuttgart library. We followed the manufacturer’s protocol for multiplex sequencing. Raw overlapping forward and reverse reads were merged and filtered for quality^51^ and mapped to the human reference genome (hg19/GRCh37/1000Genomes) using the Burrows-Wheeler Aligner (BWA)^52^ (SI2). For deeper sequencing, UDG-treated libraries of Loschbour were sequenced on 3 Illumina HiSeq 2000 lanes with 50-bp single-end reads, 8 Illumina HiSeq 2000 lanes of 100-bp paired-end reads and 8 Illumina HiSeq 2500 lanes of 101-bp paired-end reads. The UDG-treated library for Stuttgart was sequenced on 8 HiSeq 2000 lanes and 101-bp paired-end reads. The UDG-treated libraries for Motala were sequenced on 8 HiSeq 2000 lanes of 100-bp paired-end reads, with 4 lanes each for two pools (one of 3 individuals and one of 4 individuals). We also sequenced an additional 8 HiSeq 2000 lanes for Motala12, the Motala sample with the highest percentage of endogenous human DNA.

### Enrichment of mitochondrial DNA and sequencing

Non-UDG-treated libraries of Loschbour and all Motala samples were enriched for human mitochondrial DNA using a bead-based capture approach with present-day human DNA as bait^53^ to test for DNA preservation and mtDNA contamination. UDG-treatment was omitted in order to allow characterization of damage patterns typical for ancient DNA^8^. The captured libraries were sequenced on an llumina Genome Analyzer IIx platform with 2 × 76 + 7 cycles and the resulting reads were merged and quality filtered^51^. The sequences were mapped to the Reconstructed Sapiens Reference Sequence, RSRS^54^, using a custom iterative mapping assembler, MIA^55^ (SI4).

### Contamination estimates

We assessed if the sequences had the characteristics of authentic ancient DNA using four approaches. First we searched for evidence of contamination by determining whether the sequences mapping to the mitochondrial genome were consistent with deriving from more than one individual^55,56^. Second, for the high-coverage Loschbour and Stuttgart genomes, we used a maximum-likelihood-based estimate of autosomal contamination that uses variation at sites that are fixed in the 1000 Genomes data to estimate error, heterozygosity and contamination^57^ simultaneously. Third, we estimated contamination based on the rate of polymorphic sites on the X chromosome of the male Loschbour individual^58^ (SI3) Fourth, we analyzed non-UDG treated reads mapping to the RSRS to search for aDNA-typical damage patterns resulting in C→T changes at the 5'-end of the molecule^8^ (SI3).

### Phylogenetic analysis of the mitochondrial genomes

All nine complete mitochondrial genomes that fulfilled the criteria of authenticity were assigned to haplogroups using Haplofind^59^. A Maximum Parsimony tree including present day humans and previously published ancient mtDNA sequences was generated with MEGA^60^. The effect of branch shortening due to a lower number of substitutions in ancient lineages was studied by calculating the nucleotide edit distance to the root for all haplogroup R sequences (SI4).

### Sex Determination and Y-chromosome Analysis

We assessed the sex of all sequenced individuals by using the ratio of (chrY) to (chrY+chrX) aligned reads^10^. We downloaded a list of Y-chromosome SNPs curated by the International Society of Genetic Genealogy (ISOGG, http://www.isogg.org) v. 9.22 (accessed Feb. 18, 2014) and determined the state of the ancient individuals at positions where a single allele was observed and MAPQ≥30. We excluded C/G or A/T SNPs due to uncertainty about the polarity of the mutation in the database. The ancient individuals were assigned haplogroups based on their derived state (SI5). We also used BEAST v1.7.51^61^ to assess the phylogenetic position of Loschbour using 623 males from around the world with 2,799 variant sites across 500kb of non-recombining Y-chromosome sequence^62^ (SI5).

### Estimation of Neandertal admixture

We estimate Neandertal admixture in ancient individuals with the *f_4_-*ratio or *S*-statistic^5,63,64^ *αˆ* = *f*_4_ (*Altai, Denisova; Test, Yoruba)/ f*_4_(*Altai, Denisova; Vindija, Yoruba*) which uses whole genome data from Altai, a high coverage (52×) Neanderthal genome sequence^35^, Denisova, a high coverage sequence^49^ from another archaic human population (31×), and Vindija, a low coverage (1.3×) Neanderthal genome from a mixture of three Neanderthal individuals from Vindija Cave in Croatia^63^.

### Inference of demographic history and inbreeding

We used the Pairwise Sequentially Markovian Coalescent (PSMC)^65^ to infer the size of the ancestral population of Stuttgart and Loschbour. This analysis requires high quality diploid genotype calls and cannot be performed in the low-coverage Motala samples. To determine whether the low effective population size inferred for Loschbour is due to recent inbreeding, we plotted the time-to-most-recent common ancestor (TMRCA) along each of chr1-22 to detect runs of low TMRCA.

### Analysis of segmental duplications and copy number variants

We built read-depth based copy number maps for the Loschbour, Stuttgart and Motala12 genomes in addition to the Denisova and Altai Neanderthal genome and 25 deeply sequenced modern genomes^35^ (SI7). We built these maps by aligning reads, subdivided into their non-overlapping 36-bp constituents, against the reference genome using the mrsFAST aligner^66^, and renormalizing read-depth for local GC content. We estimated copy numbers in windows of 500 unmasked base pairs slid at 100 bp intervals across the genome. We called copy number variants using a scale space filter algorithm. We genotyped variants of interest and compared the genotypes to those from individuals sequenced as part of the 1000 Genomes Project^67^.

### Phenotypic inference

We inferred likely phenotypes (SI8) by analyzing DNA polymorphism data in the VCF format^68^ using VCFtools (http://vcftoools.sourceforge.net/). For the Loschbour and Stuttgart individuals, we included data from sites not flagged as LowQuality, with genotype quality (GQ) of ≥30, and SNP quality (QUAL) of ≥50. For Motala12, which is of lower coverage, we included sites having at least 2× coverage and passed visual inspection of the local alignment using samtools tview (http://samtools.sourceforge.net)^69^

### Human Origins dataset curation

The Human Origins array consists of 14 panels of SNPs for which the ascertainment is well known^5,70^. All population genetics analysis were carried out on a set of 594,924 autosomal SNPs, after restricting to sites that had >90% completeness across 7 different batches of sequencing, and that had >97.5% concordance with at least one of two subsets of samples for which whole genome sequencing data was also available. The total dataset consists of 2,722 individuals, which we filtered to 2,345 individuals (203 populations) after removing outlier individuals or relatives based on visual inspection of PCA plots^15,71^ or model-based clustering analysis^14^. Whole genome amplified (WGA) individuals were not used in analysis, except for a Saami individual who we forced in because of the special interest of this population for Northeastern European population history (Extended Data Fig. 7).

### ADMIXTURE analysis

We merged all Human Origins genotype data with whole genome sequencing data from Loschbour, Stuttgart, MA1, Motala12, Motala_merge, and LaBrana. We then thinned the resulting dataset to remove SNPs in linkage-disequilibrium with PLINK 1.07^72^, using a window size of 200 SNPs advanced by 25 SNPs and an r^2^ threshold of 0.4. We ran ADMIXTURE 1.23^14,73^ for 100 replicates with different starting random seeds, default 5-fold cross-validation, and varying the number of ancestral populations K between 2 and 20. We assessed clustering quality using CLUMPP^74^. We used the ADMIXTURE results to identify a set of 59 “West Eurasian” (European/Near Eastern) populations based on values of a “West Eurasian” ancestral population at K=3 (SI9). We also identified 15 populations for use as “non-West Eurasian outgroups” based on their having at least 10 individuals and no evidence of European or Near Eastern admixture at K=11, the lowest K for which Near Eastern/European-maximized ancestral populations appeared consistently across all 100 replicates.

### Principal Components Analysis

We used *smartpca^15^* (version: 10210) from EIGENSOFT^71,75^ 5.0.1 to carry out Principal Components Analysis (PCA) (SI10). We performed PCA on a subset on individuals and then projected others using the *lsqproject: YES* option that gives an unbiased inference of the position of samples even in the presence of missing data (especially important for ancient DNA).

### *f_3_*-statistics

We use the *f_3_*-statistic^5^ 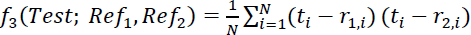, where *t_i_*, *r*_1_,_i_ and *r*_2,i_ are the allele frequencies for the *i*^th^ SNP in populations *Test, Ref_1_*, *Ref_2_*, respectively, to determine if there is evidence that the *Test* population is derived from admixture of populations related to *Ref*_1_ and *Ref*_2_ (SI11). A significantly negative statistic provides unambiguous evidence of mixture in the *Test* population^5^. We allow *Ref_1_* and *Ref_2_* to be any Human Origins population with 4 or more individuals, or Loschbour, Stuttgart, MA1, Motala12, LaBrana. We assess significance of the *f_3_*-statistics using a block jackknife^21^ and a block size of 5cM. We report significance as the number of standard errors by which the statistic differs from zero (Z-score). We also perform an analysis in which we constrain the reference populations to be (i) EEF (Stuttgart) and WHG (Loschbour or LaBrana), (ii) EEF and a Near Eastern population, (iii) EEF and ANE (MA1), or (iv) any two present-day populations, and compute a Z_diff_ score between the lowest *f_3_*-statistic observed in the dataset, and the *f_3_-*statistic observed for the specified pair.

### *f_4_*-statistics

We analyze *f_4_*-statistics^5^ of the form 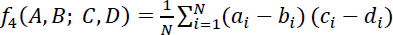 to assess if populations A, B are consistent with forming a clade in an unrooted tree with respect to C, D. If they form a clade, the allele frequency differences between the two pairs should be uncorrelated and the statistic has an expected value of 0. We set the outgroup *D* to be a sub-Saharan African population or Chimpanzee. We systematically tried all possible combinations of the ancient samples or 15 “non-West Eurasian outgroups” identified by ADMIXTURE analysis as A, B, C to determine their genetic affinities (SI14). Setting A as a present-day test population and B as either Stuttgart or BedouinB, we documented relatedness to C=(Loschbour or MA1) or C=(MA1 and Karitiana) or C=(MA1 or Han) (Extended Data Figs. 4, 5, 7). Setting C as a test population and (A, B) a pair from (Loschbour, Stuttgart, MA1) we documented differential relatedness to ancient populations (Extended Data Fig. 6). We computed *D*-statistics^63^ using transversion polymorphisms in whole genome sequence data^35^ to confirm robustness to ascertainment and ancient DNA damage (Extended Data Table 2).

### Minimum number of source populations for Europeans

We used *qpWave*^22,23^ to study the minimum number of source populations for a designated set of Europeans (SI12). We use *f_4_*-statistics of the form *X(l, r) = f_4_(l_0_, l; r_0_, r)* where *l_0_*,*r_0_* are arbitrarily chosen “base” populations, and *l*, *r* are other populations from two sets *L* and *R* respectively. If *X(l, r)* has rank *r* and there were *n* waves of immigration into *R* with no back-migration from *R* to *L*, then *r+1 ≤ n*. We set *L* to include *Stuttgart, Loschbour, MA1, Onge, Karitiana, Mbuti* and *R* to include 23 modern European populations who fit the model of SI14 and had admixture proportions within the interval [0,1] for the method with minimal modeling assumptions (SI17).

### Admixture proportions for Stuttgart in the absence of a Near Eastern ancient genome

We used Loschbour and BedouinB as surrogates for “Unknown hunter-gatherer” and Near Eastern (NE) farmer populations that contributed to Stuttgart (SI13). Ancient Near Eastern ancestry in Stuttgart is estimated by the *f_4_*-ratio^5,18^ *f_4_(Outgroup, X; Loschbour, Stuttgart) / f_4_(Outgroup, X; Loschbour, NE)*. A complication is that BedouinB is a mixture of NE and African ancestry. We therefore subtracted^23^ the effects of African ancestry using estimates of the BedouinB African admixture proportion from ADMIXTURE (SI9) or ALDER^76^.

### Admixture graph modeling

We used ADMIXTUREGRAPH^5^ (version 3110) to model population relationships between Loschbour, Stuttgart, Onge, and Karitiana using Mbuti as an African outgroup. We assessed model fit using a block jackknife of differences between estimated and fitted *f*-statistics for the set of included populations (we expressed the fit as a Z score). We determined that a model failed if |Z|>3 for at least one *f*-statistic. A basic tree model failed and we manually amended the model to test all possible models with a single admixture event, which also failed. Further manual amendment to include 2 admixture events resulted in 8 successful models, only one of which could be amended to also fit MA1 as an additional constraint. We successfully fit both the Iceman and LaBrana into this model as simple clades and Motala12 as a 2-way mixture. We also fit present-day West Eurasians as clades, 2-way mixtures, or 3-way mixtures in this basic model, achieving a successful fit for a larger number of European populations (n=26) as 3-way mixtures. We estimated the individual admixture proportions from the fitted model parameters. To test if fitted parameters for different populations are consistent with each other, we jointly fit all pairs of populations *A* and *B* by modifying ADMIXTUREGRAPH to add a large constant (10,000) to the variance term *f_3_(A_0_, A, B).* By doing this, we can safely ignore recent gene flow within Europe that affects statistics that include both *A* and *B*.

### Ancestry estimates from *f_4_*-ratios

We estimate EEF ancestry using the *f_4_*-ratio^5,18^ *f_4_(Mbuti, Onge; Loschbour, European)* / *f_4_(Mbuti, Onge; Loschbour, Stuttgart),* which produces consistent results with ADMIXTUREGRAPH (SI14). We use *f_4_(Stuttgart, Loschbour; Onge MA1)* / *f_4_(Mbuti, MA1; Onge, Loschbour)* to estimate Basal Eurasian admixture into Stuttgart. We use *f_4_(Stuttgart, Loschbour; Onge Karitiana)* / *f_4_(Stuttgart, Loschbour; Onge MA1)* to estimate ANE mixture in Karitiana (Fig. 2B). We use *f_4_(Test, Stuttgart; Karitiana, Onge)* / *f_4_(MA1, Stuttgart; Karitiana, Onge)* to lower bound ANE mixture into North Caucasian populations.

### MixMapper analysis

We carried out *MixMapper* 2.0^4^ analysis, a semi-supervised admixture graph fitting technique. First, we infer a scaffold tree of populations without strong evidence of mixture relative to each other (Mbuti, Onge, Loschbour and MA1). We do not include European populations in the scaffold as all had significantly negative *f_3_*-statistics indicating admixture. We then ran *MixMapper* to infer the relatedness of the other ancient and present-day samples, fitting them onto the scaffold as 2- or 3-way mixtures. The uncertainty in all parameter estimates is measured by block bootstrap resampling of the SNP set (100 replicates with 50 blocks).

### *TreeMix* analysis

We applied *TreeMix*^34^ to Loschbour, Stuttgart, Motala12, and MA1^6^, LaBrana^12^ and the Iceman^19^, along with the present-day samples of Karitiana, Onge and Mbuti. We restricted the analysis to 265,521 Human Origins array sites after excluding any SNPs where there were no-calls in any of the studied individuals. The tree was rooted with Mbuti and standard errors were estimated using blocks of 500 SNPs. We repeated the analysis on whole-genome sequence data, rooting with Chimp and replacing Onge with Dai since we did not have Onge whole genome sequence data^35^. We varied the number of migration events (*m*) between 0 and 5.

### Inferring admixture proportions with minimal modeling assumptions

We devised a method to infer ancestry proportions from three ancestral populations (EEF, WHG, and ANE) without strong phylogenetic assumptions (SI17). We rely on 15 “non-West Eurasian” outgroups and study *f_4_(European, Stuttgart; O_1_, O_2_)* which equals *αβ f_4_(Loschbour, Stuttgart; O_1_, O_2_)* + *α(1-β) f_4_(MA1, Stuttgart; O_1_, O_2_)* if *European* has 1-*a* ancestry from EEF and *β*, 1-*β* ancestry from WHG and ANE respectively. This defines a system of 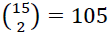 equations with unknowns *αβ*, *α*(1-*β*), which we solve with least squares implemented in the function *lsfit* in *R* to obtain estimates of *α* and *β*. We repeated this computation 22 times dropping one chromosome at a time^26^ to obtain block jackknife^21^ estimates of the ancestry proportions and standard errors, with block size equal to the number of SNPs per chromosome. We assessed consistency of the inferred admixture proportions with those derived from the ADMIXTUREGRAPH model based on the number of standard errors between the two (Extended Data Table 1).

### Haplotype-based analyses

We used RefinedIBD from BEAGLE 4^77^ with the settings *ibdtrim*=20 and *ibdwindow*=25 to study IBD sharing between Loschbour and Stuttgart and populations from the POPRES dataset^42^. We kept all IBD tracts spanning at least 0.5 centimorgans (cM) and with a LOD score >3 (SI18). We also used ChromoPainter^37^ to study haplotype sharing between Loschbour and Stuttgart and present-day West Eurasian populations (SI19). We identified 495,357 SNPs that were complete in all individuals and phased the data using Beagle 4^77^ with parameters *phase-its*=50 and *impute-its*=10. We did not keep sites with missing data to avoid imputing modern alleles into the ancient individuals. We combined ChromoPainter output for chromosomes 1-22 using ChromoCombine^37^. We carried out a PCA of the co-ancestry matrix using fineSTRUCTURE^37^.

## References

1. Bramanti, B. et al. Genetic discontinuity between local hunter-gatherers and Central Europe’s first farmers. Science 326, 137–140, (2009).

2. Skoglund, P. et al. Origins and genetic legacy of Neolithic farmers and hunter-gatherers in Europe. Science 336, 466–469, (2012).

3. Haak, W. et al. Ancient DNA from European early Neolithic farmers reveals their Near Eastern affinities. PLoS Biol. 8, e1000536, (2010).

4. Lipson, M. et al. Efficient moment-based inference of admixture parameters and sources of gene flow. Mol. Biol. Evol. 30, 1788–1802, (2013).

5. Patterson, N. et al. Ancient admixture in human history. Genetics 192, 1065–1093, (2012).

6. Raghavan, M. et al. Upper Palaeolithic Siberian genome reveals dual ancestry of Native Americans. Nature 505, 87–91, (2014).

7. Krause, J. et al. A complete mtDNA genome of an early modern human from Kostenki, Russia. Curr. Biol. 20, 231–236, (2010).

8. Sawyer, S., Krause, J., Guschanski, K., Savolainen, V. & Pääbo, S. Temporal patterns of nucleotide misincorporations and DNA fragmentation in ancient DNA. PLoS ONE 7, e34131, (2012).

9. Haak, W. et al. Ancient DNA from the first European farmers in 7500-Year-old Neolithic sites. Science 310, 1016–1018, (2005).

10. Skoglund, P., Storå, J., Götherström, A. & Jakobsson, M. Accurate sex identification of ancient human remains using DNA shotgun sequencing. J. Archaeol. Sci. 40, 4477–4482, (2013).

11. Soares, P. et al. The Archaeogenetics of Europe. Curr. Biol. 20, R174–R183, (2010).

12. Olalde, I. et al. Derived immune and ancestral pigmentation alleles in a 7,000-year-old Mesolithic European. Nature 507, 225–228, (2014).

13. Perry, G. H. et al. Diet and the evolution of human amylase gene copy number variation. Nat. Genet. 39, 1256–1260, (2007).

14. Alexander, D. H., Novembre, J. & Lange, K. Fast model-based estimation of ancestry in unrelated individuals. Genome Res. 19, 1655–1664, (2009).

15. Patterson, N., Price, A. L. & Reich, D. Population structure and eigenanalysis. PLoS Genet. 2, e190, (2006).

16. Lao, O. et al. Correlation between genetic and geographic structure in Europe. Curr. Biol. 18, 1241–1248, (2008).

17. Novembre, J. et al. Genes mirror geography within Europe. Nature 456, 98–101, (2008).

18. Reich, D., Thangaraj, K., Patterson, N., Price, A. L. & Singh, L. Reconstructing Indian population history. Nature 461, 489–494, (2009).

19. Keller, A. et al. New insights into the Tyrolean Iceman's origin and phenotype as inferred by whole-genome sequencing. Nat. Commun. 3, 698, (2012).

20. Novembre, J. & Stephens, M. Interpreting principal component analyses of spatial population genetic variation. Nat. Genet. 40, 646–649, (2008).

21. Busing, F. T. A., Meijer, E. & Leeden, R. Delete-m Jackknife for Unequal m. Statistics and Computing 9, 3–8, (1999).

22. Moorjani, P. et al. Genetic evidence for recent population mixture in India. Am. J. Hum. Genet. 93, 422–438, (2013).

23. Reich, D. et al. Reconstructing Native American population history. Nature 488, 370–374, (2012).

24. Botigué, L. R. et al. Gene flow from North Africa contributes to differential human genetic diversity in southern Europe. Proc. Natl. Acad. Sci. USA 110, 11791–11796, (2013).

25. Cerezo, M. et al. Reconstructing ancient mitochondrial DNA links between Africa and Europe. Genome Res. 22, 821–826, (2012).

26. Moorjani, P. et al. The history of African gene flow into southern Europeans, Levantines, and Jews. PLoS Genet. 7, e1001373, (2011).

27. Thangaraj, K. et al. Reconstructing the origin of Andaman Islanders. Science 308, 996–996, (2005).

28. Fu, Q. et al. DNA analysis of an early modern human from Tianyuan Cave, China. Proc. Natl. Acad. Sci. USA 110, 2223–2227, (2013).

29. Bar-Yosef, O. The chronology of the Middle Paleolithic of the Levant. 39–56 (New York: Plenum Press, 1998).

30. Armitage, S. J. et al. The southern route “Out of Africa”: evidence for an early expansion of modern humans into Arabia. Science 331, 453–456, (2011).

31. Rose, J. I. et al. The Nubian Complex of Dhofar, Oman: an African middle stone age industry in Southern Arabia. PLoS ONE 6, e28239, (2011).

32. Haber, M. et al. Genome-Wide Diversity in the Levant Reveals Recent Structuring by Culture. PLoS Genet 9, e1003316, (2013).

33. Brace, C. L. et al. The questionable contribution of the Neolithic and the Bronze Age to European craniofacial form. Proc. Natl. Acad. Sci. U. S. A. 103, 242–247, (2006).

34. Pickrell, J. K. & Pritchard, J. K. Inference of population splits and mixtures from genome-wide Allele frequency data. PLoS Genet. 8, e1002967, (2012).

35. Prufer, K. et al. The complete genome sequence of a Neanderthal from the Altai Mountains. Nature 505, 43–49, (2014).

36. Ralph, P. & Coop, G. The geography of recent genetic ancestry across Europe. PLoS Biol. 11, e1001555, (2013).

37. Lawson, D. J., Hellenthal, G., Myers, S. & Falush, D. Inference of Population Structure using Dense Haplotype Data. PLoS Genet. 8, e1002453, (2012).

38. Hellenthal, G. et al. A genetic atlas of human admixture history. Science 343, 747–751, (2014).

39. Bellwood, P. First Farmers: The Origins of Agricultural Societies. (Wiley-Blackwell, 2004).

40. Brandt, G. et al. Ancient DNA reveals key stages in the formation of central European mitochondrial genetic diversity. Science 342, 257–261, (2013).

## References

41. McKenna, A., et al. The Genome Analysis Toolkit: a MapReduce framework for analyzing next-generation DNA sequencing data. Genome Res. 20, 1297–1303, (2010).

42. Nelson, M. R. et al. The Population Reference Sample, POPRES: a resource for population, disease, and pharmacological genetics research. Am. J. Hum. Genet. 83, 347–358, (2008).

43. Delsate, D., Guinet, J.-M. & Saverwyns, S. De l'ocre sur le crâne mésolithique (haplogroupe U5a) de Reuland-Loschbour (Grand-Duché de Luxembourg) ? Bull. Soc. Préhist. Luxembourgeoise 31, 7–30, (2009).

44. Rohland, N. & Hofreiter, M. Ancient DNA extraction from bones and teeth. Nat. Protocols 2, 1756–1762, (2007).

45. Dabney, J. et al. Complete mitochondrial genome sequence of a Middle Pleistocene cave bear reconstructed from ultrashort DNA fragments. Proceedings of the National Academy of Sciences 110, 15758–15763, (2013).

46. Stäuble, H. S. f. V.-u. F. d. U. F. Häuser und absolute Datierung der Ältesten Bandkeramik. (Habelt, 2005).

47. Yang, D. Y., Eng, B., Waye, J. S., Dudar, J. C. & Saunders, S. R. Improved DNA extraction from ancient bones using silica-based spin columns. Am. J. Phys. Anthropol. 105, 539–543, (1998).

48. Meyer, M. & Kircher, M. Illumina sequencing library preparation for highly multiplexed target capture and sequencing. Cold Spring Harb. Protoc. 2010, pdb prot5448, (2010).

49. Meyer, M. et al. A High-Coverage Genome Sequence from an Archaic Denisovan Individual. Science 338, 222–226, (2012).

50. Briggs, A. W. et al. Removal of deaminated cytosines and detection of in vivo methylation in ancient DNA. Nucleic Acids Res. 38, e87–e87, (2010).

51. Kircher, M. in *Methods Mol*. Biol. Vol. 840 Methods in Molecular Biology 197–228 (2012).

52. Li, H. & Durbin, R. Fast and accurate short read alignment with Burrows–Wheeler transform. Bioinformatics 25, 1754–1760, (2009).

53. Maricic, T., Whitten, M. & Pääbo, S. Multiplexed DNA Sequence Capture of Mitochondrial Genomes Using PCR Products. PLoS ONE 5, e14004, (2010).

54. Behar, Doron M. et al. A Copernican Reassessment of the Human Mitochondrial DNA Tree from its Root. Am. J. Hum. Genet. 90, 675–684, (2012).

55. Green, R. E. et al. A Complete Neandertal Mitochondrial Genome Sequence Determined by High-Throughput Sequencing. Cell 134, 416–426, (2008).

56. Fu, Q. et al. A Revised Timescale for Human Evolution Based on Ancient Mitochondrial Genomes. Curr. Biol. 23, 553–559, (2013).

57. Fu, Q. & al., e. *(in preparation),* (2014).

58. Rasmussen, M. et al. An Aboriginal Australian Genome Reveals Separate Human Dispersals into Asia. Science 334, 94–98, (2011).

59. Vianello, D., et al. HAPLOFIND: a new method for high-throughput mtDNA haplogroup assignment. Hum. Mutat. 34, 1189–1194, (2013).

60. Tamura, K., et al. MEGA5: Molecular Evolutionary Genetics Analysis using Maximum Likelihood, Evolutionary Distance, and Maximum Parsimony Methods. Mol. Biol. Evol. 28, 2731–2739, (2011).

61. Drummond, A. & Rambaut, A. BEAST: Bayesian evolutionary analysis by sampling trees. BMC Evol. Biol. 7, 214, (2007).

62. Lippold, S. et al. Human paternal and maternal demographic histories: insights from high-resolution Y chromosome and mtDNA sequences. bioRxiv, doi: 10.1101/001792, (2014).

63. Green, R. E. et al. A Draft Sequence of the Neandertal Genome. Science 328, 710–722, (2010).

64. Reich, D. et al. Genetic history of an archaic hominin group from Denisova Cave in Siberia. Nature 468, 1053–1060, (2010).

65. Li, H. & Durbin, R. Inference of human population history from individual whole-genome sequences. Nature 475, 493–496, (2011).

66. Hach, F. et al. mrsFAST: a cache-oblivious algorithm for short-read mapping. Nat. Meth. 7, 576–577, (2010).

67. An integrated map of genetic variation from 1,092 human genomes. Nature 491, 56–65, (2012).

68. Danecek, P. et al. The variant call format and VCFtools. Bioinformatics 27, 2156–2158, (2011).

69. Li, H. The sequence alignment/map (SAM) format and SAMtools. Bioinformatics. 25, 2078–2079, (2009).

70. Keinan, A., Mullikin, J. C., Patterson, N. & Reich, D. Measurement of the human allele frequency spectrum demonstrates greater genetic drift in East Asians than in Europeans. Nat Genet 39, 1251–1255, (2007).

71. Price, A. L. et al. Principal components analysis corrects for stratification in genome-wide association studies. Nat. Genet. 38, 904–909, (2006).

72. Purcell, S. et al. PLINK: a tool set for whole-genome association and population-based linkage analyses. Am. J. Hum. Genet. 81, 559–575, (2007).

73. Alexander, D. & Lange, K. Enhancements to the ADMIXTURE algorithm for individual ancestry estimation. BMC Bioinformatics 12, 246, (2011).

74. Jakobsson, M. & Rosenberg, N. A. CLUMPP: a cluster matching and permutation program for dealing with label switching and multimodality in analysis of population structure. Bioinformatics 23, 1801–1806, (2007).

75. Price, A. L., Zaitlen, N. A., Reich, D. & Patterson, N. New approaches to population stratification in genome-wide association studies. Nat. Rev. Genet. 11, 459–463, (2010).

76. Loh, P.-R. et al. Inferring Admixture Histories of Human Populations Using Linkage Disequilibrium. Genetics 193, 1233–1254, (2013).

77. Browning, B. L. & Browning, S. R. Improving the Accuracy and Efficiency of Identity-by-Descent Detection in Population Data. Genetics 194, 459–471, (2013).

